# HELZ2: a new, interferon-regulated, human 3’-5’ exoribonuclease of the RNB family is expressed from a non-canonical initiation codon

**DOI:** 10.1101/2023.02.22.529493

**Authors:** Eric Huntzinger, Jordan Sinteff, Bastien Morlet, Bertrand Seraphin

**Author notes:** 77500 Chelles, France (Jordan Sinteff).

## Abstract

Proteins containing a RNB domain, originally identified in *E. coli* RNase II, are widely present throughout the tree of life. Many RNB proteins are endowed with 3’-5’ exoribonucleolytic activity but some have lost catalytic function during evolution. Database searches identified a new RNB domain containing protein in human: HELZ2. Analysis of genomic and expression data with evolutionary information suggested that the human HELZ2 protein is produced from an unforeseen non-canonical initiation codon in Hominidae. This unusual property was confirmed experimentally, extending the human protein by 247 residues. Human HELZ2 was further shown to be an active ribonuclease despite the substitution of a key residue in its catalytic center. HELZ2 harbors also two RNA helicase domains and several zinc-fingers and its expression is induced by interferon treatment. We demonstrate that HELZ2 is able to degrade structured RNAs through the coordinated ATP-dependent displacement of duplex RNA mediated by its RNA helicase domains and its 3’-5’ ribonucleolytic action. The expression characteristics and biochemical properties of HELZ2 support a role for this factor in response to viruses and/or mobile elements.

## Introduction

During their lifecycle, RNA molecules interact with numerous proteins that allow their synthesis, maturation, localization, translation and finally their degradation. Among the many factors interacting with RNA, ribonucleases play a unique role in participating both to the initial maturation of newly made RNAs and in their final degradation. Proteins belonging to numerous protein families are endowed with ribonuclease activity targeting cellular RNAs (1). Some of these factors are also mobilized by cells to defend themselves against pathogens such as RNA viruses, but may in some cases be diverted by the latter to facilitate their propagation.

A major ribonuclease family is defined by the presence of a RNB domain that catalyzes 3’-5’ hydrolytic RNA degradation. Pioneering members of this group, RNase II and RNase R, were identified in *E. coli* (2). If related proteins are widely distributed in bacteria, they are also found in eukaryotes as well as in some archaea and viruses. In eukaryotes, a highly conserved RNB domain protein is the catalytic subunit of a decameric complex named the exosome that constitutes the major cellular 3’-5’ exonuclease in both the nucleus and the cytoplasm (3–5). If the exosome appears highly conserved, some other RNB proteins are only present in some specific clades of the eukaryotic tree. Hence, in human, 3 RNB-domain containing proteins have been characterized: DIS3 and DIS3L1 that are exosome subunits (6, 7) and DIS3L2 that acts as an independent enzyme targeting uridylated RNAs (8). If most proteins with a RNB domain are active nucleases, some have lost a catalytic activity during evolution and are indirectly involved in RNA metabolism (9).

Database searches reveal the presence of another RNB-domain containing protein encoded by the human genome: HELZ2 (Helicase with Zinc Finger 2, also known as PDIP-1, PRIC285). Interestingly, expression of HELZ2 has been shown to be induced by interferon in mammalian cells (10), a property conserved in fish cells (11) as well as zebrafish embryo (12). In agreement, mimicking a viral infection by transfecting RNA or double-stranded DNA in mouse embryonic fibroblasts also induces HELZ2 mRNA expression (13). Analysis in chicken cells demonstrated that HELZ2 is controlled by the MDA5 receptor that senses double stranded RNA (14). However, not all RNA viruses induce HELZ2 expression in every cell type, since infection of Saos-2 human osteosarcoma cells by Zika virus has no effect (15). Consistent with regulation of HELZ2 by interferon and pattern recognition receptors, a study reported that it mediates the inhibition of Dengue virus infection in human hepatic cells induced by IFN-α (16). In line with this observation, downregulation of HELZ2 expression reduces the antiviral effects of IFN-α against hepatitis C virus (HCV) infection (17). More recently, HELZ2 was found among the genes whose expression is upregulated in cells following SARS-CoV-2 infection (18, 19). A CRISPR screen suggested further that HELZ2 may have a proviral action for SARS-Cov2 (20). Beyond human systems, control of virus by HELZ2 extends to birds as inhibition of HELZ2 facilitated infection by Duck Tembusu virus (21). A part from viruses, HELZ2 was also reported to inhibit retrotransposition of human LINE1 (22). A link of HELZ2 with immune response was also established by population genetic: associations of the HELZ2 locus to an autoimmune disease locus in B cells and to an autoimmune liver disease called primary biliary cholangitis were reported (23, 24). Finally, *Helz2*-knockout mice were shown to be resistant to high-fat diet-induced obesity, glucose intolerance, and hepatosteatosis (25). These phenotypes can be explained by a central leptin resistance and by the increase in leptin receptor mRNA in the liver of the mutant mice, leading to a decrease in lipogenesis as well as an increase in fatty acid oxidation.

Altogether, these data clearly link HELZ2 to pathogens in particular RNA viruses and possibly other physiological functions. To study the potential contribution of HELZ2 to these processes, we decided to characterize the molecular activity of the human protein. Detailed analyses uncovered that human HELZ2 is longer than the sequence currently listed in databases as it is translated from a non-canonical translation codon. Analysis of the full-length HELZ2 sequence revealed that it contains various domains including several copies of two types of Zn fingers and 2 helicase domains in addition to the RNB domain. Biochemical analyses demonstrate that, despite the presence of a non-canonical residue in its RNB domain active site, HELZ2 is an active 3’-5’ exonuclease. We show further that the RNA helicase domains synergize with the RNB domain to allow degradation of structured RNA, a feature that may be important to target viruses or structured RNAs.

## Material and Methods

### Construction of eukaryotic expression vectors

Sequences encoding HELZ2β and HELZ2α were amplified respectively from p3XFLAG-CMV_7.1_ HELZ2α and p3XFLAG-CMV_7.1_ HELZ2β (kind gift from Dr. T. Tomaru), and inserted between the EcoRI-SalI sites of pEGFP-C1 vector (Clontech) giving pBS5988 and pBS5528. For the cloning of the sequence encoding the N-terminal extension, a fragment of 897 nucleotides, covering the extension and the first 756 nucleotides of HELZ2β, overlapping the first exon, was amplified from genomic DNA and inserted into EcoRI-ClaI sites into peGFPC1-HELZ2β to give pBS6283. Sequence encoding human Dis3L1 was amplified from HEK293 cDNA preparation and then inserted between XhoI and SalI sites of pEGFP-C1 vector giving pBS6465. Subsequent modifications of these vectors were made by mutagenesis PCR. Original clones and derivatives were verified by Sanger sequencing.

For luciferases assays, complementary oligos containing the 5’UTR of HELZ2 mRNA (nts 1 to 151) and the first 35 nts of firefly luciferase were annealed and extended by PCR. The resulting product was then digested and inserted between HindIII-KasI sites of pGL3 vector (Promega) giving pBS6286. Subsequent modifications of this vector were made by mutagenesis PCR. Original clones and derivatives were verified by Sanger sequencing. Oligonucleotides and plasmids used for this study are listed in Supplementary Table 1 and 2 respectively.

### Cell culture and transfections

HEK293 cells (ATCC CRL-1573) were maintained in DMEM medium containing 4.5 g/l glucose, GlutaMAX, 10% fetal calf serum and 40 μg/ml gentamicin. They were transfected with Lipofectamine^2000^ transfection reagent (ThermoFischer) according to the manufacturer’s recommendation and processed 24 to 48 hours post-transfection.

HeLa cells were maintained in DMEM medium containing 1 g/l glucose, 10% fetal calf serum and 40 μg/ml gentamicin. For interferon β stimulation, cells were seeded in 6 well plates and then mock-treated or treated with 5000 U/mL of human recombinant IFNβ (R&D systems, 8499-IF/CF) for 24 hours.

### Luciferase Assay

HEK293 cells were seeded in 12 well plates and then co-transfected with 500 ng of firefly luciferase and 250 ng of renilla luciferase vectors. After 24 hours, cells are washed with PBS1X and then lysed with 250 μl of 1X Passive Lysis Buffer (Promega). Firefly and renilla luminescences were then monitored with Dual-Glo Luciferase Assay kit (Promega) in a Berthold luminometer. Firefly values were normalized with renilla values to control for transfection efficiency.

### Microscopy

HEK293 cells were grown on coverslips and transfected with pEGFPC1-HELZ2. After 48 hours, cells were fixed with 4% paraformaldehyde that was then neutralized by incubation with 0,125M glycine followed by 3 washes with PBS1X. Cells are then incubated for 30 sec with DAPI diluted at 1/10000. After 3 more washes, coverslips with cells were deposited on a glass slide together with one drop of mounting medium (ProLong Gold antifade reagent from Invitrogen). Microscopic images were taken with a Leica DM4000 B with an objective HC PL APO 100×1.40 oil CS2 and equipped with a Hamamatsu ORCA-Flash 4.0 LT C11440 camera.

### Western blotting

Western blotting was performed using standard procedures and blots visualized with Amersham Imager 600 or 800 (GE Healthcare). GFP fusion proteins were revealed with mouse monoclonal antibody anti-GFP (JL-8, Clontech) used at 1/2000 dilution. Anti-mouse IgG HRP linked antibody (Cell Signaling Technology, 7076) was used as secondary antibody and revelation was performed with the Luminata Crescendo Western HRP Substrate (Millipore).

### Oligonucleotides and RNA labelling

RNA and DNA oligonucleotides (100 pmol) were 5’ end-labeled using T4 PNK (ThermoFischer) and γ-P^32^ ATP according to manufacturer instructions. Reactions were stopped by addition of 5 μl of 0,25M EDTA, supplemented with 2X RNA loading dye (ThermoFischer) and loaded on a 6% acryl-urea 8M gel for purification. After migration, bands corresponding to radiolabeled oligonucleotides were identified by autoradiography, excised and eluted overnight at 4°C in elution buffer (20 mM Tris-HCl pH 7.5, 0,25 M sodium acetate, 1 mM EDTA, 0,25% SDS). The next day, supernatants were recovered, phenol-chloroform extracted and precipitated. The resulting pellets were resuspended in 11 μl of ddH2O. RNA and DNA oligonucleotides used for the assays are listed in Supplementary Table 3.

### Production of RNA-RNA or RNA-DNA duplexes

5’ end-labeled RNA was incubated with 3-fold excess of complementary cold RNA in annealing buffer (60 mM KCl, 6 mM Hepes pH 7,5, 0,2 mM MgCl_2_). After 2 min incubation at 90°C, the mixture was allowed to cool slowly to room temperature. Samples were then loaded on a 6% native acrylamide gel in 1X TBE and fractionated by electrophoresis. After migration, an autoradiography was performed to locate bands of interest and those were cut and eluted overnight at 4°C in elution buffer (20 mM Tris-HCl pH 7.5, 0,25 M sodium acetate, 1 mM EDTA, 0,25% SDS). The next day, supernatants were recovered, phenol-chloroform extracted, and precipitated. The resulting pellets were resuspended in 11 μl of ddH2O.

### GFP-tagged protein immunoprecipitation for biochemical assay

Immunoprecipitation experiments were performed using GFP-Trap magnetic agarose beads (Chromotek) as recommended by the manufacturer. Specifically, 48 hours after transfection cells grown in 15 cm dishes were lysed for 20 min on ice in 500 μl lysis buffer (10 mM Tris-HCl pH 8.0, 150 mM NaCl, 0.5% Igepal CA-630, 1mM DTT) supplemented with protease inhibitors (1X Complete Protease Inhibitor Cocktail EDTA-free, Roche). After centrifugation at 20,000 g for 20 min at 4°C, supernatants were recovered and protein concentration was determined with Protein Assay Dye Reagent Concentrate (Bio-Rad). Per assay, 1 mg of cell lysates were then incubated with 5 μl GFP-Trap paramagnetic beads for 1 hour at 4 °C mixing by rotation. After recovery, beads were washed three times with 1 ml wash buffer (10 mM Tris-HCl pH 8.0, 500 mM NaCl, 0.5% Igepal CA-630, 1mM DTT). During the last wash, 10% of the beads were recovered for western blot analysis. The rest of the beads were aliquoted in separated tubes are then resuspended in 45 μl of reaction buffer (10 mM Tris-HCl pH 8, 150 mM NaCl, 5 mM MgCl_2_ and 1 mM DTT) and kept on ice until further use.

### RNase assay

To proteins bound to GFP-Trap beads (see above), 5 μl of a mix containing radiolabeled RNA and radiolabeled DNA diluted in reaction buffer (10 mM Tris-HCl pH 8, 150 mM NaCl, 5 mM MgCl_2_ and 1 mM DTT) were added. Aliquots of 5 μl were taken at different time points and mixed with 1 μl of stop reaction buffer (1% SDS, 0,1 M EDTA) and 6 μl of 2X RNA loading dye (ThermoFischer). Samples were then incubated for 2 min at 90°C and then one third was loaded on a 20% acryl-urea 8M gel for separation. After the migration, gels were exposed to an imaging plate (Fuji) for 6 hours and then revealed using a Typhoon FLA-9500. Bands were quantified using ImageQuant.

### ATPase assay

To proteins bound to GFP-Trap beads (see above), 5 μl of a mix containing reaction buffer, α-P^32^ ATP (25 pmol) were added. DNA (100 pmol) or RNA (100 pmol) were included as indicated. Aliquots of 10 μl were taken from reactions after 2 hours and mixed with 1 μl of stop reaction buffer (1% SDS, 0,1 M EDTA). One microliter of each sample was then loaded on a thin-layer chromatography plate (PEI cellulose F, Merck) and resolved with KH_2_PO_4_ 0,75M buffer. In parallel, ATP and ADP were loaded as migration controls. ADP was produced by incubating radioactive ATP with cold ADP and nucleotide di-phosphate kinase (NDK, N2635-100UN, Sigma-Aldrich) for 30 min at 25°C. After the migration, dried TLC plates were exposed to an imaging plate (Fuji) for 2 hours and then revealed using a Typhoon FLA-9500. Signals were quantified using ImageQuant.

### Assays with guanabenz acetate

RNase and ATPase assays were performed as described in the presence or absence of guanabenz acetate (sc-203590, Santa Cruz).

### Sample preparation for mass spectrometry analyses

After 24 hours of stimulation with interferon or mock treatment, cells were recovered and then lysed in RIPA buffer (50 mM Tris-HCl pH7,4, 150 mM NaCl, 1% Igepal, 0,5% sodium deoxycholate, 0,1% SDS). Bradford assay was used to determine protein concentrations and equal amounts of extracts were further analyzed by mass spectrometry. The protein mixtures were precipitated with TCA 20% overnight at 4°C and centrifuged at 14,000 rpm for 10 min at 4°C. Protein pellets were washed twice with 1 mL cold acetone and air dried. The protein extracts were solubilized in urea 2 M, reduced with 5 mM TCEP for 30 min and alkylated with 10 mM iodoacetamide for 30 min in the dark. Enzymatic digestion was performed at 37°C and overnight with 500 ng trypsin (Promega, Charbonnieres les Bains, France). Peptide mixtures were then desalted on C18 spin-column and dried in a Speed-Vacuum.

### LC-MS/MS Analysis

Samples were analyzed using an Ultimate 3000 nano-RSLC coupled in line, via the NanoFlex-electrospray ionization source, with the Orbitrap Exploris 480 mass-spectrometer (Thermo Fisher Scientific, Bremen, Germany) equipped with a FAIMS (high Field Asymmetric Ion Mobility Spectrometry) module. Peptide mixtures were injected in 0.1% TFA on a C18 Acclaim PepMap100 trap-column (75 μm ID x 2 cm, 3 μm, 100Å, Thermo Fisher Scientific) for 3 min at 6 μL/min with 2% ACN, 0.1% FA in H2O and then separated on a BioZen peptide XB-C18 nano-column (75 μm ID x 25 cm, 2.6 μm, Phenomenex) at 350 nl/min and 45°C with a 60 min linear gradient from 9% to 30% buffer B (A: 0.1% FA in H2O / B: 80% ACN, 0.1% FA in H2O), regeneration at 9% B. Spray voltage were set to 2.1 kV and heated capillary temperature at 280°C.

The Orbitrap Exploris 480 was associated with the FAIMS module, set to -45 V as Compensation Voltage (CV). A cycle time of 1.2 second was chosen. For the full MS1 in DDA mode, the resolution was set to 120,000 at m/z 200 and with a mass range set to 330-1200. The full MS AGC target was 300% with a Max IT set to 100ms. For the fragment spectra in MS2, AGC target value was 100% (Standard) with a resolution of 30,000 and the maximum Injection Time set to Auto mode. Intensity threshold was set at 5E3. Isolation width was set at 2 m/z and normalized collision energy was set at 30%. All spectra were acquired in centroid mode using positive polarity. Default settings were used for FAIMS with voltage applied as described previously, and with a total carrier gas flow set to 4.2 L/min.

### Mass spectrometry data analysis

Proteins were identified by database searching using SequestHT (Thermo Fisher Scientific) with Proteome Discoverer 2.5 software (PD2.5, Thermo Fisher Scientific) on human FASTA database downloaded from UniProt (reviewed, release 2022_10_27, 20607 entries, https://www.uniprot.org/). Precursor and fragment mass tolerances were set at 7 ppm and 0.02 Da respectively, and up to 2 missed cleavages were allowed. For all the data, Oxidation (M, +15.995 Da) was set as variable modification, and Carbamidomethylation (C, + 57.021 Da) as fixed modification. Peptides and proteins were filtered with a false discovery rate (FDR) at 1%. Label-free quantification was based on the extracted ion chromatography intensity of the peptides and realized with Perseus 1.6.15.0 (Max Planck Institute of Biochemistry). All samples were measured in biological triplicates. The measured extracted ion chromatogram (XIC) intensities were normalized based on median intensities of the entire dataset to correct minor loading differences. For statistical tests, not detectable intensity values were treated with an imputation method, where the missing values were replaced by random values similar to the 10% of the lowest intensity values present in the entire dataset. Unpaired two tailed T-test, assuming equal variance, were performed on obtained log2 XIC intensities.

## Results

### Human HELZ2 protein translation initiates with a non-canonical start codon

HELZ2 (also named PRIC285 or PDIP1) was initially identified by two independent groups in experiments aiming to identify novel partners for the DNA binding domains of PPARα and PPARγ (Peroxisomal Proliferator Activated Receptor alpha and gamma) (26, 27). A first isoform, named HELZ2α and corresponding to a protein of 231kDa, was identified by cDNA sequencing in the original study (26) (Fig. 1A). The second study predicted a larger, N-terminally extended, isoform of 295kDa termed HELZ2β (27) (Fig. 1A). While the two isoforms could be encoded by alternatively spliced mRNAs, 5’ RACE experiments performed by the second group failed to amplify the 5’ end of the cDNA encoding the α isoform suggesting that it was at best of low abundance (27). This conclusion is further supported by the results of RNase protection assays performed by the second team (27) and the mapping of transcription initiation sites that failed to confirm the start site of the HELZ2α mRNA (28). Moreover, a single mRNA corresponding in size to the one encoding the β isoform, and significantly longer than the one predicted to encode HELZ2α, was detected in Northern blots by the first group. Further, the predicted mRNA encoding HELZ2α contains an upstream out of frame AUG codon and a predicted 3’ splice site (26). Altogether, available data retrospectively support the existence of a single mRNA with the α isoform being probably not relevant as deduced from an incompletely spliced pre-mRNA.

**Figure 1:**
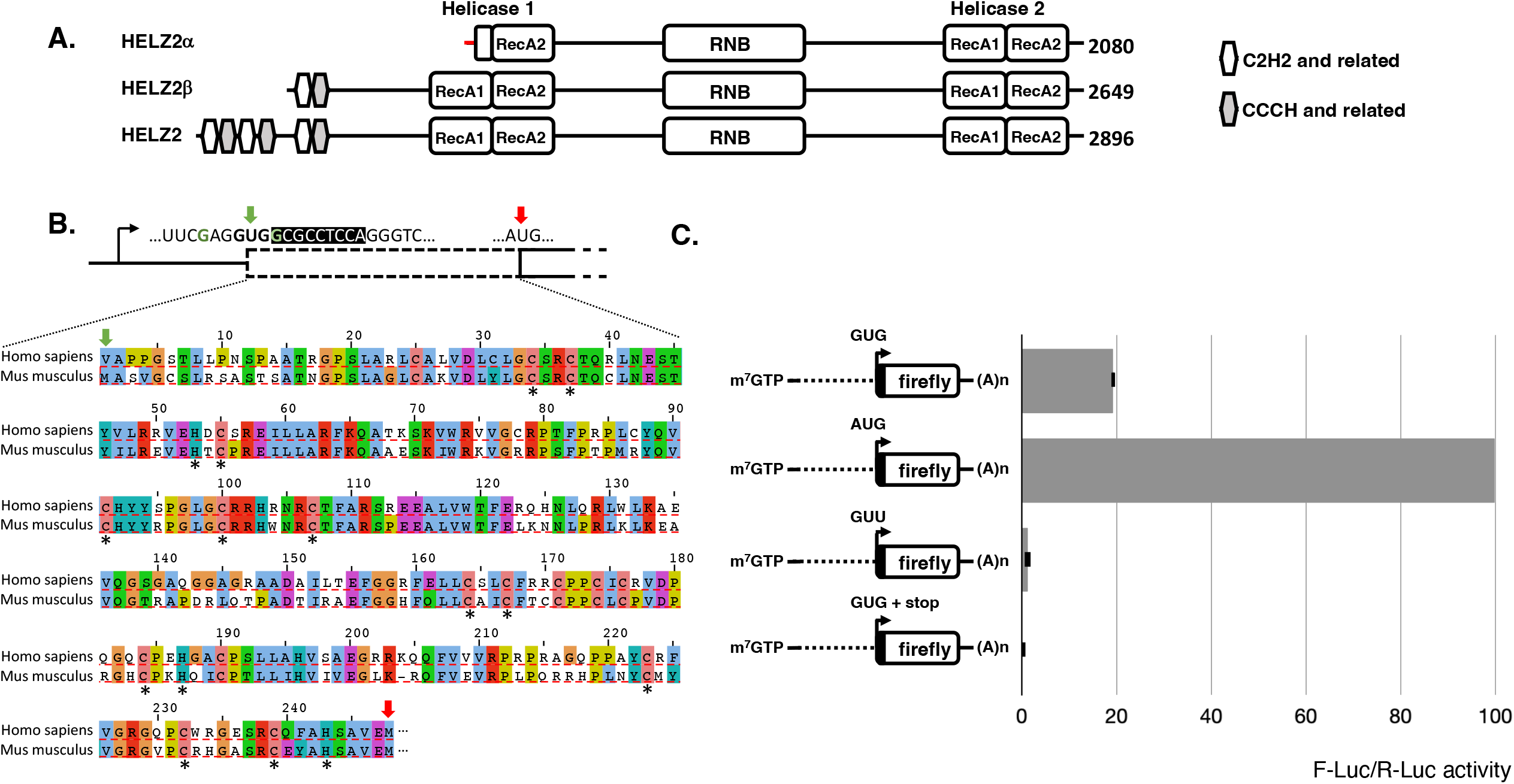
Human HELZ2 translation is initiated by a non-canonical initiation codon. (A) Domain organization of human HELZ2 and its previously described isoforms. The alternative sequence present at the N-terminus of the putative HELZ2α isoform that truncate the RecA1 domain of Helicase 1 is indicted in red. (B) Close-up view of human HELZ2 mRNA 5’ end organisation and sequence alignment of the N-terminal region of mouse HELZ2 (Uniprot: E9QAM5) and the *in silico* translated sequence of human HELZ2 N-terminal extension. Top: black arrow: transcription start site of human HELZ2 mRNA. Continuous line: 5’UTR. Dashed box: human HELZ2 N-terminal extension. Above are indicated the non-canonical GUG initiation codon (green arrow) and the canonical initiation codon of HELZ2β (red arrow). The context surrounding the GUG codon is provided with nucleotides favourable for translation initiation at the non-canonical initiation codon in green. Nucleotides following the GUG codon with black background are included in the reporters used in Panel C. Below the schematic organization, an alignment of the N-terminal region of mouse HELZ2 and the *in silico* translated sequence of human HELZ2 N-terminal extension from the amino acid encoded by the GUG codon to the previously described initiating methionine of the HELZ2β isoform. Note that is the absence of experimental data it is unclear whether the GUG start codon is decoded as M, V or both. The sequence alignment is coloured for amino acid conservation using Clustal W and asterisks placed below the sequence indicate C and H residues of the conserved additional Zn-fingers motifs. (C) Luciferase assay to evaluate HELZ2 translation initiation. Schemes of the reporters used are shown on the left. Dashed line: HELZ2 5’UTR. Black arrow: initiation codon or mutant thereof. Black box: 9 nucleotides from HELZ2 sequence following the initiating codon. Open box: luciferase coding sequence. On the right: firefly luciferase activity normalized to co-transfected renilla luciferase for each construct.

Intriguingly, the full-length cDNA contains a long 5’UTR of 893 residues (Fig. 1B) that contains 4 out of frame AUG codons (Supplementary Fig. 1). Moreover, the predicted human β isoform protein is significantly shorter than homologs in many other mammals. For example, the predicted HELZ2 protein is 2649 amino acids long in human versus 2947 for its mouse counterpart, with the main difference resulting from the absence of sequence similar to the N-terminal 247 residue of the mouse protein. Comparison of the mouse N-terminal extension with sequences encoded by the human genome indicates that a very similar sequence (65% identity) is encoded in frame and upstream of the β isoform AUG codon (Fig. 1B). This, together with the presence of a GUG codon at position 152 in the human cDNA aligning with the mouse AUG codon, suggested that HELZ2 translation in human is initiated from a non-canonical start codon. The presence of a favorable context for translation initiation (a purine at position -3 and guanosine at position +4 (29)) around the GUG codon further supported this hypothesis (Fig. 1B). However, we couldn’t rule out the possibility that human HELZ2 translation initiates at another non-canonical codon such as CUG at cDNA position 140 also in a favorable context. Analysis of published ribosome profiling data supported the translation of the sequence upstream of the β isoform initiation codon (30) (Supplementary Fig. 2). Moreover, the conservation of the unusual organization of the HELZ2 coding sequence, including the GUG codon present at position 152 in the human cDNA, in all Hominidae species argued that it is biologically relevant.

To validate the capacity of the 5’ extremity of the human HELZ2 mRNA to promote translation initiation at a non-canonical codon, we constructed a reporter in which the sequence starting at the HELZ2 mRNA transcription initiation site, extending to the GUG and including the following 9 nucleotides were fused upstream of, and in-frame with, the coding sequence of firefly luciferase. As reference, we used the same construct in which the GUG codon has been substituted by an AUG codon. HEK293 cells were transfected with the resulting plasmids together with a plasmid expressing renilla luciferase to normalize transfection efficiencies. Results indicate the GUG construct is producing a significant level of luciferase, albeit 5,2 fold reduced level compared to the AUG construct (Fig. 1C). To ascertain that the signal observed is due to translation at the GUG codon, and not to any downstream initiation event, two additional reporters were constructed. In the first one, the GUG codon at position 152 was mutated into a GUU, and in the second one, a stop codon was introduced 3 residues after the GUG codon (Fig. 1C). The GUU construct produced only traces of luciferase about 20-fold less than the GUG construct, possibly owing to inefficient translation initiation at other sites (e.g., CUG at position 140). Introducing a stop four codons downstream of the GUG codon reduced firefly luciferase levels to background (roughly 250-fold reduction compared to the GUG construct) indicating that translation emanates exclusively from the HELZ2 sequence present in our construct. Altogether, these results indicate that the GUG codon is used for translation initiation at about 20% the level of a AUG codon in the same context (Fig. 1C).

To establish whether HELZ2 translation initiates *in vivo* upstream of the AUG codon currently annotated as the start of the β isoform, we performed mass spectrometry experiments. Interferon-responsive HeLa cells were used to prepare total proteins that were analyzed by tandem mass spectrometry using as reference the human proteome present in the UNIPROT database completed by sequence predicted extension. Triplicate samples from control or interferon treated cells were prepared. Neither the HELZ2β isoform nor the extension were detected in cells growing in the absence of interferon. As expected, interferon treatment induced expression of many proteins (Fig. 2A). Those included HELZ2 and, importantly, peptides belonging to both the HELZ2β sequence and the extension were reproducibly detected after interferon stimulation (Fig. 2B). This result indicates that translation initiates *in vivo* upstream of the AUG codon currently annotated as the start of the β isoform. Interestingly, the relative numbers of Peptide Spectrum Match (PSM) for the extension and β isoform were nearly proportional to the corresponding peptide lengths (Fig. 2B), suggesting that they were expressed at similar levels. While this is only indicative, it suggests that *in vivo* produced HELZ2 includes mostly, if not always, the upstream extension.

Altogether, our data indicates that HELZ2 expression is unusual as its translation initiates from a hitherto unrecognized non-canonical start codon.

**Figure 2:**
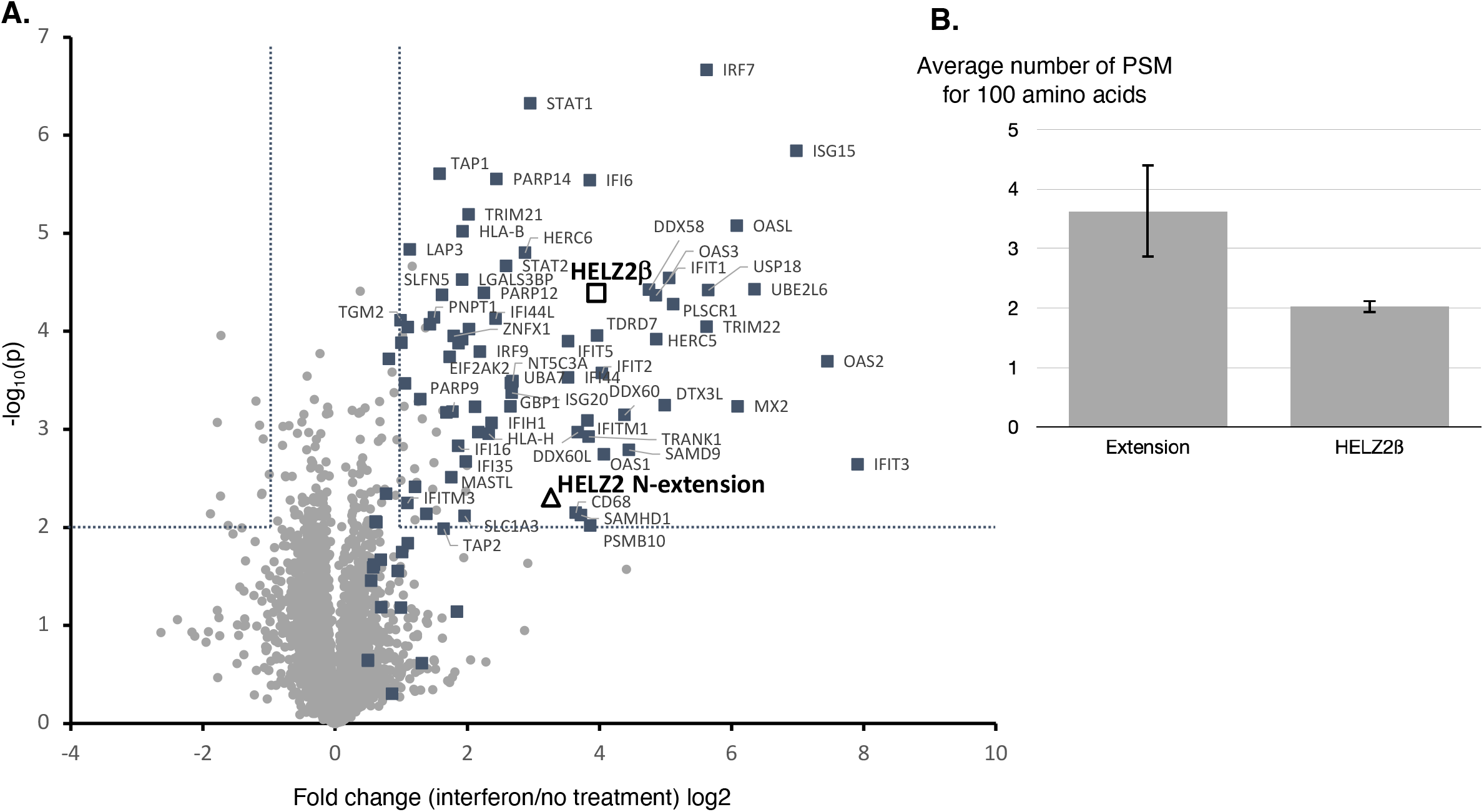
Mass spectrometry analysis of protein extracts from mock- or IFNβ treated HeLa cells. (A) Volcano-plot showing the log2 fold change versus -log_10_(p-value) between mock- or interferon treated HeLa cells. Analyses were performed in triplicate and a total of 4828 proteins were quantified by mass spectrometry. Filled or open squares indicate human proteins that whose expression at mRNA or protein levels have been reported to be induced at least 3 fold by interferon according to the Interferome Database (58). The names of some of these factors are indicated. Open square and triangle represent measurement for peptide corresponding to the HELZ2β isoform and the HELZ2 extension, respectively. All other proteins are indicated by grey dots. Dashed lines delimit regions of p-value higher lower than 0,01 and up- or down-regulation of at least 2 fold. (B) Graphical representation of the number of PSM per 100 amino acids for the N-terminal extension sequence of HELZ2 and the HELZ2β isoform.

### Domain organization of human HELZ2

Database searches and multiple sequence alignments reveal that human HELZ2 has a modular structure (Fig. 1A). Full-length human HELZ2 contains 6 Zn-finger or Zn-finger like domains at its N-terminus. These appear to be of two types with some departing from the corresponding consensus. Yet, their characteristics, the alternating positioning of the two types, and the conservation of this arrangement in related proteins support these assignments (Supplementary Fig. 3). The 1^st^, 3^rd^ and 5^th^ Zn-fingers are related to C2H2 domains that have been found in a number of proteins involved in RNA metabolism (31, 32). Some of them were proposed to mediate interaction with double-stranded RNA. The alternating Zn fingers are of the CCCH family known to bind single-stranded RNA in a sequence specific manner (33). This organization, including the presence of deviant members and additional conserved C and H amino acids suggest that the Zn-fingers may cooperate to recognize complex RNA motifs. Alternatively, they could act independently, allowing HELZ2 to recognize different RNA partners in a combinatorial manner. Interestingly, 4 of these Zn fingers are encoded by the previously unnoticed N-terminal region of the human protein (Fig. 1B). Their conservation in other animals support further the translation of this region.

The N-terminal Zn finger region is followed by a RNA helicase domain, the RNB domain and a second helicase domain (Fig. 1A, Supplementary Fig. 4). Both helicase domains belong to the Upf1-like subgroup of the SF1 helicase superfamily. The 11 proteins of this subfamily mostly interact with RNA and are monomeric SF1-B 5′–3′ helicases (34). In between the two helicase domains, lies the RNB domain that, like for most proteins harboring this signature, is preceded by two cold-shock domains (CSD). The latter have been shown to guide the incoming substrate to the exoribonucleolytic catalytic site (Supplementary Fig. 4). Most RNB domains are followed by a S1 domain that also interacts with single-stranded RNA. Accordingly, the sequence of HELZ2, and corresponding 3D-structure predicted by AlphaFold, suggest the presence of a structure with some characteristics of S1 domains downstream of the RNB domain of HELZ2. Yet, direct functional and structural analyses will be necessary to validate these predictions and unravel interactions between the different modules present in HELZ2.

### The HELZ2 RNB domain of exhibits ribonuclease activity in vitro

The HELZ2 RNB domain is unusual as one of the four highly conserved aspartic acids thought to be catalytic in proteins containing an RNB domain (2, 3, 35) is replaced with an asparagine (Fig. 3A). Interestingly, the canonical aspartate is found in HELZ2 from monotremes and other vertebrates (Supplementary Fig. 4). This observation questions whether the RNB domain of human HELZ2 lost its catalytic activity during mammalian evolution. To test whether human HELZ2 is endowed with nuclease activity we turned to *in vitro* assays. As our attempts to produce recombinant protein in bacteria or in baculovirus were unsuccessful, we used HEK293 cells transfected with the plasmids encoding GFP-HELZ2 or GFP-HELZ2β for our biochemical tests. As controls we used GFP-MBP as well as derivatives of GFP-HELZ2 and GFP-HELZ2β carrying a D to N substitution at position corresponding to position 1601 of full length HELZ2 (1354 for HELZ2β) as this residue has been shown to coordinate a catalytic magnesium (35) and its mutation in other RNB family member abolish their activity (2, 3). Proteins were immunoprecipitated from extracts with GFP-Trap beads which were extensively washed before addition of 5’ radiolabeled 35 nucleotides long synthetic DNA oligonucleotide and 20 nucleotides long synthetic RNA oligonucleotide in magnesium containing buffers. The reaction was incubated at 37°C with regular shaking and aliquots were collected at different time points. In parallel, protein samples were collected in order to analyze the material present in the input and on the beads by western blot (Fig. 3B). We observed that the GFP-MBP, GFP-HELZ2, GFP-HELZ2β wild type or mutant are all produced and immunoprecipitated in similar quantities allowing assessment of their nuclease activities (Fig. 3B). We did not observe any change in the intensity of the 35 nucleotide DNA band at all incubation times, regardless of the protein used (Fig. 3C). This indicates that the HELZ2 protein and its derivatives do not have deoxyribonuclease activity. Analysis of the fate of the RNA oligonucleotide incubated in the presence of GFP-MBP reveals only background degradation after 1 hour of incubation possibly owing to traces of contaminants (Fig. 3C). In contrast, in the presence of either HELZ2 or HELZ2β, a strong decrease in the signal corresponding to the 20 nucleotides RNA was observed even at the earliest time points (only 2 minutes of incubation) (Fig. 3C, D). Concomitantly with the disappearance of the full-length RNA substrate, we observe the apparition of degradations intermediates of progressively decreasing size during the time course experiment. These observations are consistent with the exonucleolytic 3’ to 5’ degradation of the substrate typical of those catalyzed by RNB domain containing proteins. A final product of 4-5 nucleotides starts to accumulate after 8 minutes and is predominant at the two latest time points. Importantly, the observed nuclease activity is lost when proteins harboring the point mutation in the catalytic domain were used (*rnb*^*-*^, Fig. 3C, D). We compared the size of the final product with the one generated by another RNB domain containing protein, DIS3L1, by performing ribonuclease assays side-by-side. The final products were of comparable size (Supplementary Fig. 5), a result that is also with previous analyses (3). Altogether, these results demonstrate that, despite the presence of a non-canonical residue in its active site, human HELZ2 is an active ribonuclease acting through its RNB domain. Moreover, we couldn’t detect a significant difference in ribonuclease activity between the full-length HELZ2 and the previously described shorter HELZ2β derivative (Fig. 3D).

**Figure 3:**
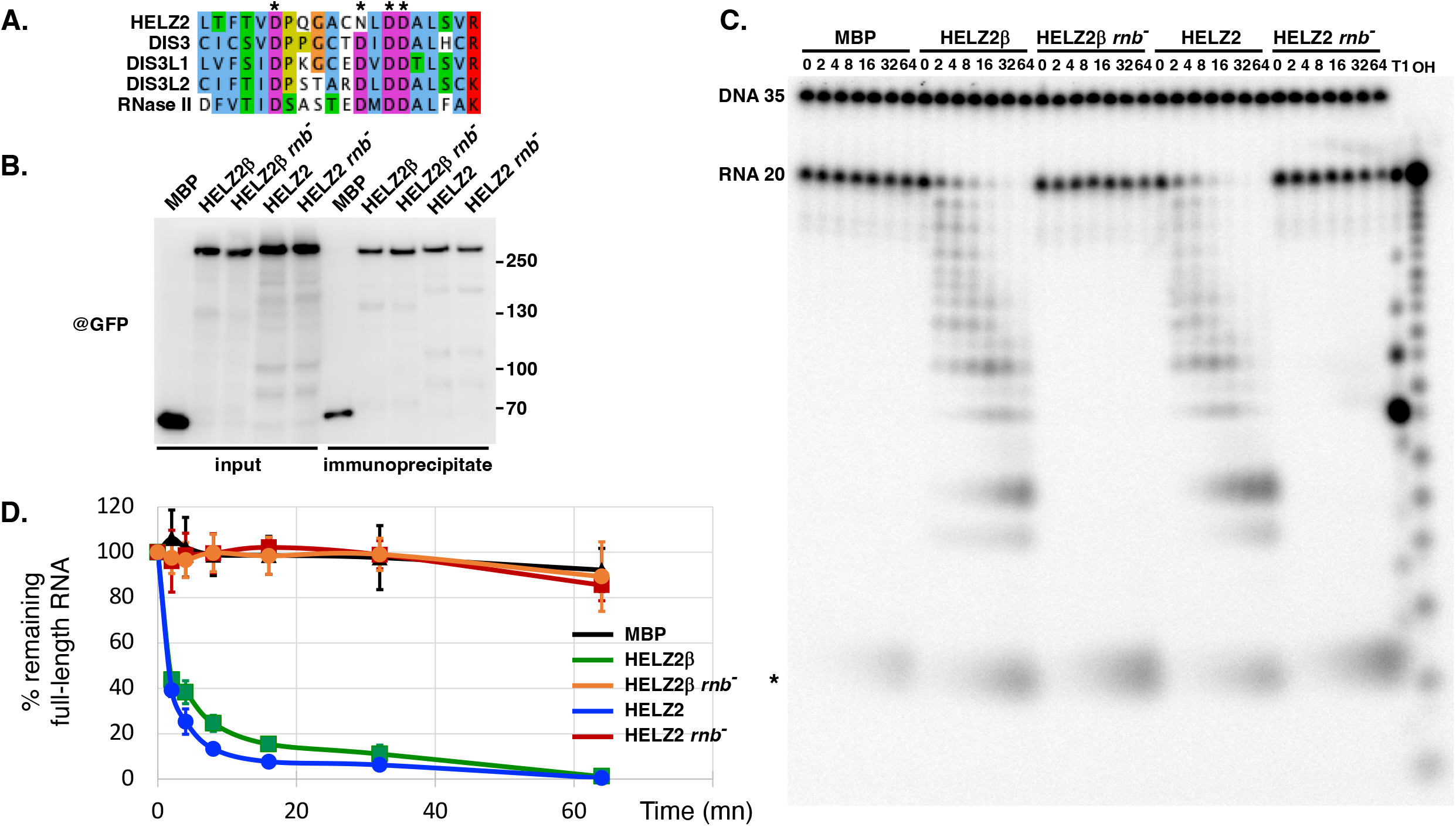
HELZ2 and HELZ2β are active ribonucleases. (A) Sequence alignment of the RNB domain catalytic region from human HELZ2 (Q9BYK8), DIS3 (Q9Y2L1), DIS3L1 (Q8TF46), DIS3L2 (Q8IYB7) and from bacterial RNaseII (P30850). Previously proposed catalytic residues are indicated by asterisks. The sequence alignment is coloured for amino acid conservation using Clustal W. (B) Western-blot using anti-GFP showing expression levels and immunoprecipitation efficiency for the proteins used in the ribonuclease activity test. (C) Representative RNA degradation assay gel. Samples from the ribonuclease activity test were fractionated on a 20% acryl-urea 8M gel. Numbers above the lanes represent incubations times in minutes. T1: RNase T1 digestion of the substrate RNA. OH: alkaline hydrolysis of the radiolabelled RNA substrate. Asterisk: non-specific product of the reaction. (D) Quantification of the bands corresponding to the full-length RNA substrate plotted against the time of the reaction in minutes with standard error (n=3).

Interestingly, one study recently reported that guanabenz acetate binds HELZ2β with high affinity and activates thereby hepatic leptin receptor expression both *in vitro* and *in vivo*, a property currently under clinical study (36, 37). Thus, we tested in vitro the degradation of RNA in presence of inhibitory concentrations of guanabenz acetate. We didn’t observe an effect of this compound on the nuclease activity of HELZ2 (Supplementary Fig. 6), suggesting that it may affect other function(s) of the protein.

### HELZ2 shows a preference for pyrimidines toward purines nucleotides

Careful examination of the degradation profiles produced by HELZ2 shows that it doesn’t progress uniformly, pausing at certain positions. Matching these points with the RNA sequence indicated that intermediates accumulate when HELZ2 encounters a guanosine at the 3’ end of the substrate RNA. To investigate more thoroughly the substrate specificity of HELZ2, we analyzed degradation of hybrid oligonucleotides composed of 10 DNA residues at their 5’ end followed by a 12-residue long 3’ oligo(A), oligo(U), oligo(C) or oligo(G) (The presence of 10 DNA residues ensured uniform labelling of the different substrates.). Time-course analysis of the degradation of these substrates using the previously described assay demonstrated a rapid degradation of the oligo(U), less efficient degradation of oligo(C), and limited degradation of oligo(A) (Fig. 3A). In contrast, the oligo(G) substrate was barely digested. Comparing the sizes of the final products with the size of a 10 nucleotide long radiolabeled DNA oligo fractionated in parallel, reveal that HELZ2 degrades the oligo(U) and oligo(C) substrates until only 2 or 3 ribonucleotides remain on the substrate. This indicates tolerance for a DNA backbone in the substrate molecule until a certain point in the path leading to the catalytic center of the protein. Altogether, these data confirm that HELZ2 is unable to degrade DNA substrate and indicate that it has a strong preference of HELZ2 for pyrimidine over purine residues for its nuclease activity.

### HELZ2 exhibits nucleic acid stimulated ATPase activity emanating from its helicase domains

The 2 helicase domains flanking the RNB domain of HELZ2 contains key amino acid involved in the binding and hydrolysis of ATP. Thus, using on-bead assay described above, we investigated whether HELZ2 is endowed with ATPase activity. The full-length GFP-HELZ2 protein was used for these reactions with GFP-MBP serving as negative control. We also introduced point mutations at residues essential positions for the binding and hydrolysis of ATP in the motif II of the two helicase domains of HELZ2: DE914/915AA and DE2607/2608AA, to ensure that activities were carried-out by HELZ2. Mutants of individual domains (hel1^-^, hel2^-^) as wells as the double mutant (hel1/2^-^) were constructed. ATPase activity was monitored following incubation with radiolabeled αATP by detecting ADP production using Thin Layer Chromatography (TLC) and autoradiography. Incubation of HELZ2 with only ATP resulted in detection of very low ATPase level after 2 hours of incubation. Addition of single-stranded nucleic acids, either RNA or DNA, to the reaction significantly increased the production of ADP (Fig. 5). As control, incubation of control GFP-MBP protein resulted in the production of only background levels of ADP. Testing the role of individual domains using mutant indicated that both domains were active and stimulated by nucleic acids. Interestingly, helicase domain 2 of HELZ2 (hel1^-^ construct) was more active than helicase domain 1 (Fig. 5). Moreover, activities of individual domains weren’t additive suggesting a possible coordination of the two helicases. We also investigated the impact of guanabenz acetate on this activity. Again, this compound didn’t block ATP hydrolysis suggesting that it doesn’t modulate the helicase function of HELZ2 (Supplementary Fig. 6).

**Figure 4:**
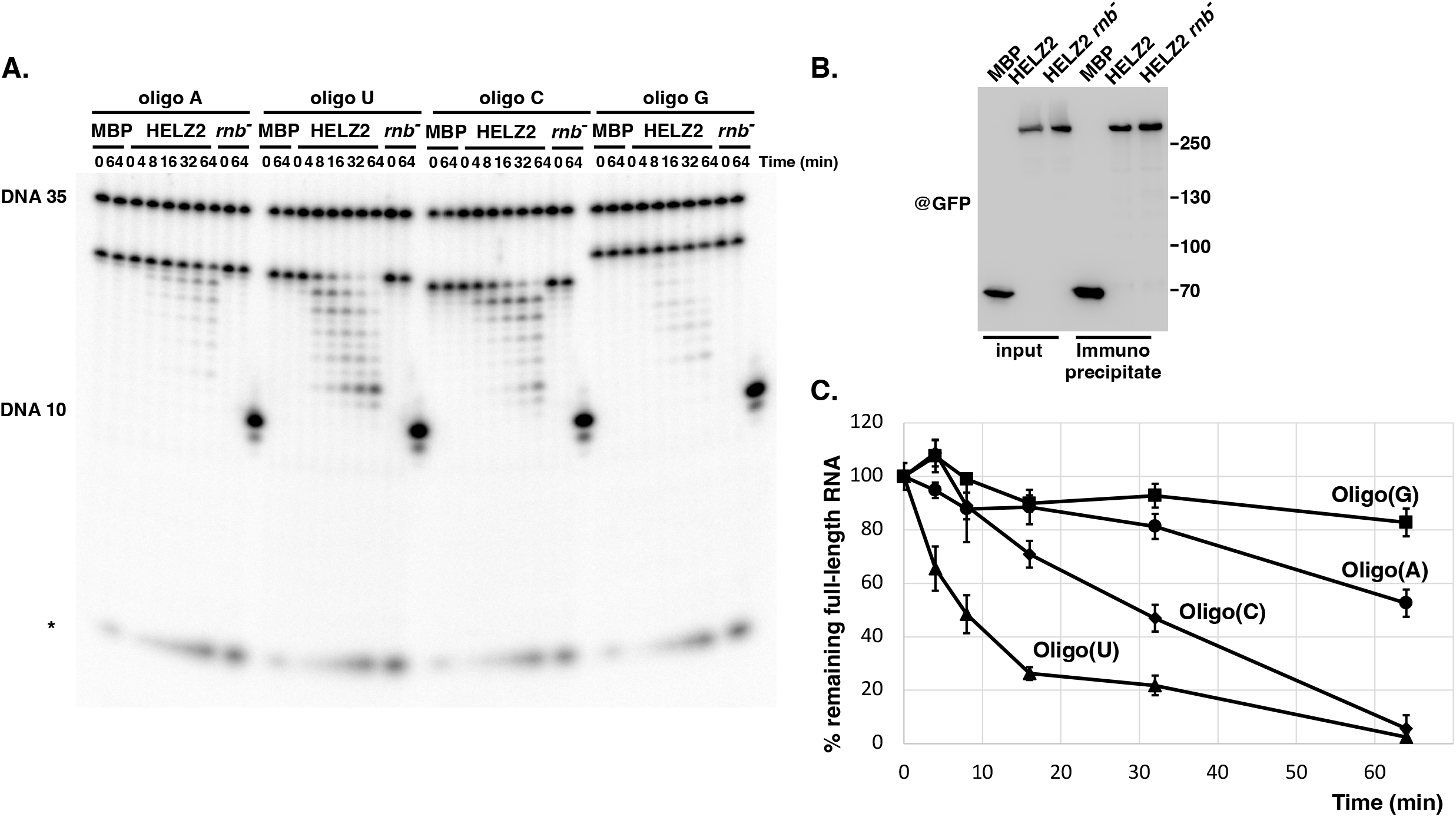
HELZ2 displays substrate preference for pyrimidines. (A) Representative RNA degradation assay gel with oligo(A), oligo(U), oligo(C) or oligo(G) substrates. Samples from the ribonuclease activity test were fractionated on a 20% acryl-urea 8M gel. Numbers above the lanes represent incubations times in minutes. A radiolabelled 10 nucleotide long DNA (DNA 10) was also loaded as reference to determine the sizes of the finals degradations products. Asterisk: non-specific product of the reaction. (B) Western-blot using anti-GFP showing expression levels and immunoprecipitation efficiency for the proteins used in the ribonuclease activity test. (C) Quantification of the bands corresponding to the full-length DNA-RNA substrates plotted against the time of the reaction in minutes with standard error (n=3).

**Figure 5:**
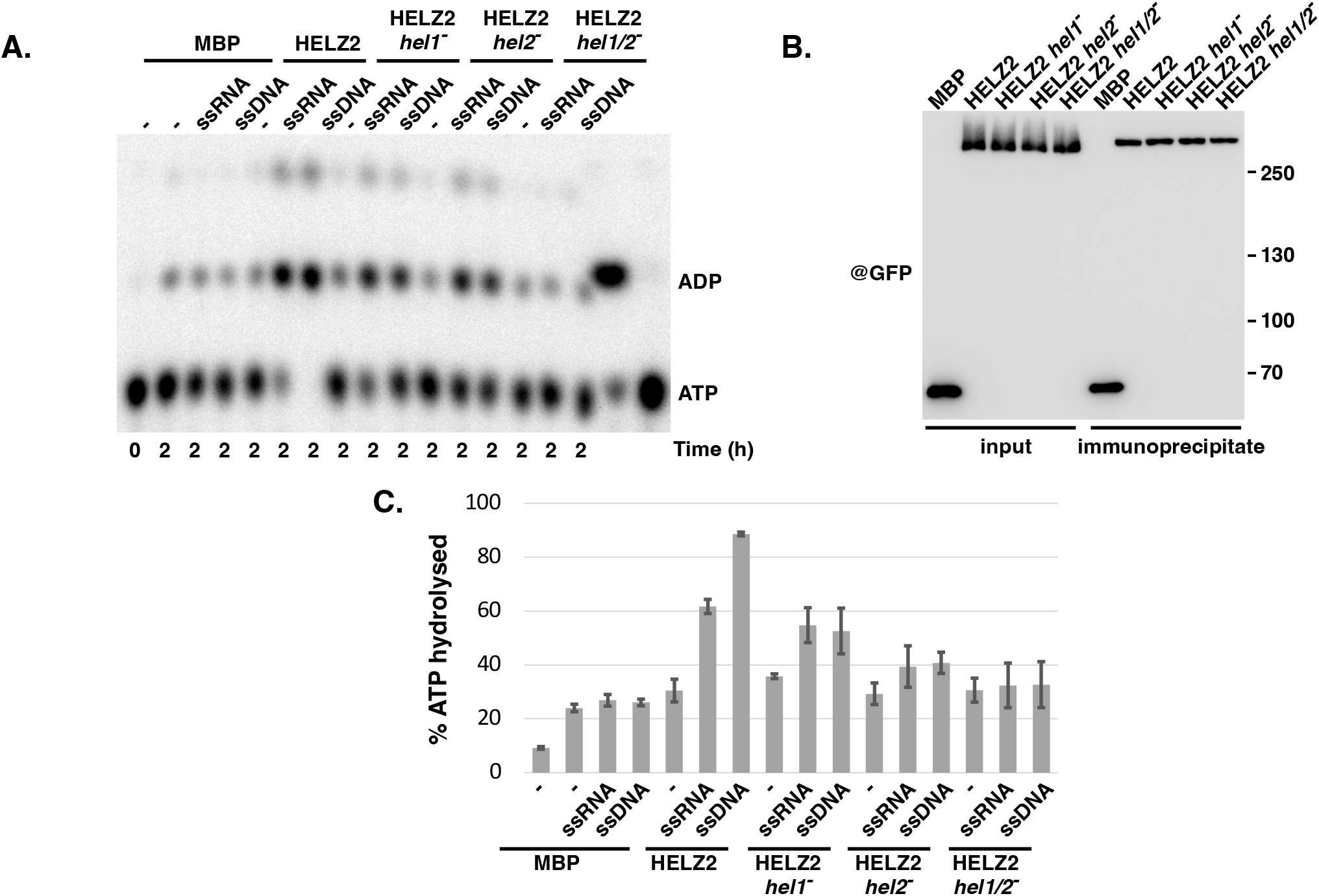
HELZ2 displays nucleic acids stimulated ATPase activity. (A) Representative ATPase assay thin layer chromatography (TLC). Samples from the ATPase activity test fractionated on a TLC and detected by autoradiography. Numbers under the lanes represent incubation times in hours. Above the lanes are indicated the nucleic acids substrates used to stimulate ATPase activity and the proteins used for the different assays. The last 2 lanes are references identify the migration of ATP and ADP, respectively. The faint signal at the top represents free phosphate. (B) Western-blot using anti-GFP showing expression levels and immunoprecipitation efficiency for the proteins used in the ATPase activity test. (C) Quantification of the percentage of hydrolysed ATP in the various conditions with standard error (n=3).

### The HELZ2 nuclease and helicase domains collaborate to degrade structured RNA

The peculiar organization of HELZ2 that include a unique combination of active helicase domains with a RNB domain may facilitate substrate to the ribonucleolytic active site. Indeed, members of the RNB family were reported to have the ability to degrade duplexes RNA substrates only as long as it contains a minimal single-stranded 3’ end (4).To test whether HELZ2 can degrade structured RNA a duplex substrate was produced by annealing a 30 nucleotide RNA with a complementary 14 nucleotide RNA, leaving 16 nucleotides single-stranded as 3’ end (Fig. 6). Degradation of this duplex by HELZ2 is stopped after the removal of a maximum of 5 residues. Addition of ATP did not allow the degradation to proceed further (Fig. 6). In control reactions, the single stranded 30 nt RNA is efficiently digested, with accumulation of numerous degradation intermediates and the presence of the final 4-5 nt long product. Interestingly, we noticed that addition of ATP stimulated single stranded RNA degradation. This enhancement was only detected when the helicase domains were active. To test the possibility that a single-stranded entry points is necessary for the unwinding, and thus degradation, of duplex RNA, we designed a substrate incorporating the 14 nt long duplex but in which the complementary strand was prolonged by a 16 nt long single stranded 5’ overhang (Fig. 6). This substrate was degraded by HELZ2 beyond the block imposed by the duplex RNA, but only when ATP was present and when the helicase domains were active (Fig. 6) Altogether, these results indicate that the helicase domains of HELZ2 collaborate with the RNB domain, giving the protein the ability to degrade structured RNA.

**Figure 6:**
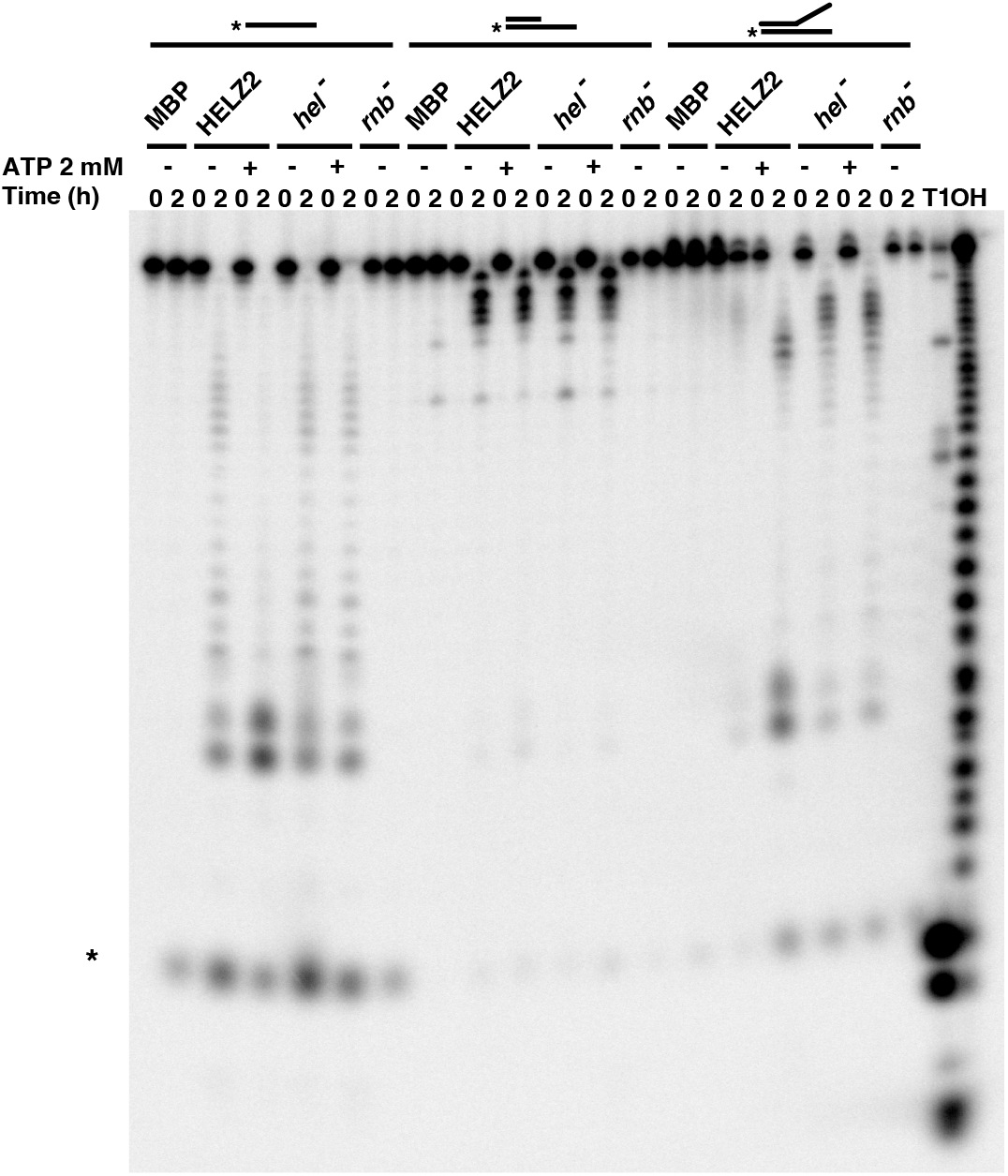
Helicase and ribonuclease activities of HELZ2 synergize for the degradation of structured RNA. Representative RNA degradation assay of duplex RNA substrates in presence or absence of ATP. Samples from the ribonuclease activity test were fractionated on a 20% acryl-urea 8M gel. Numbers above the lanes represent incubations times in hours. Above the figure is a schematic representation of the substrates and proteins used for the assay. T1: RNase T1 digestion of the substrate. OH: alkaline hydrolysis of the radiolabelled RNA substrate. Asterisk: non-specific product of the reaction.

### GFP-HELZ2 localizes mainly localized in the cytoplasm

HELZ2 was originally categorized as a nuclear protein colocalizing with PPARα (26) but more recent data suggest that HELZ2 may function in the cytoplasm to fulfill some of its functions (22). To solve this contradiction, we tested the localization of transiently expressed GFP-HELZ2 by microscopy analyses. Our results indicate that GFP-HELZ2 is predominantly a cytoplasmic protein with an overall diffuse distribution. In some cells, we noticed the presence of few bright foci that were probably enriched in GFP-HELZ2 (Fig. 7).

**Figure 7:**
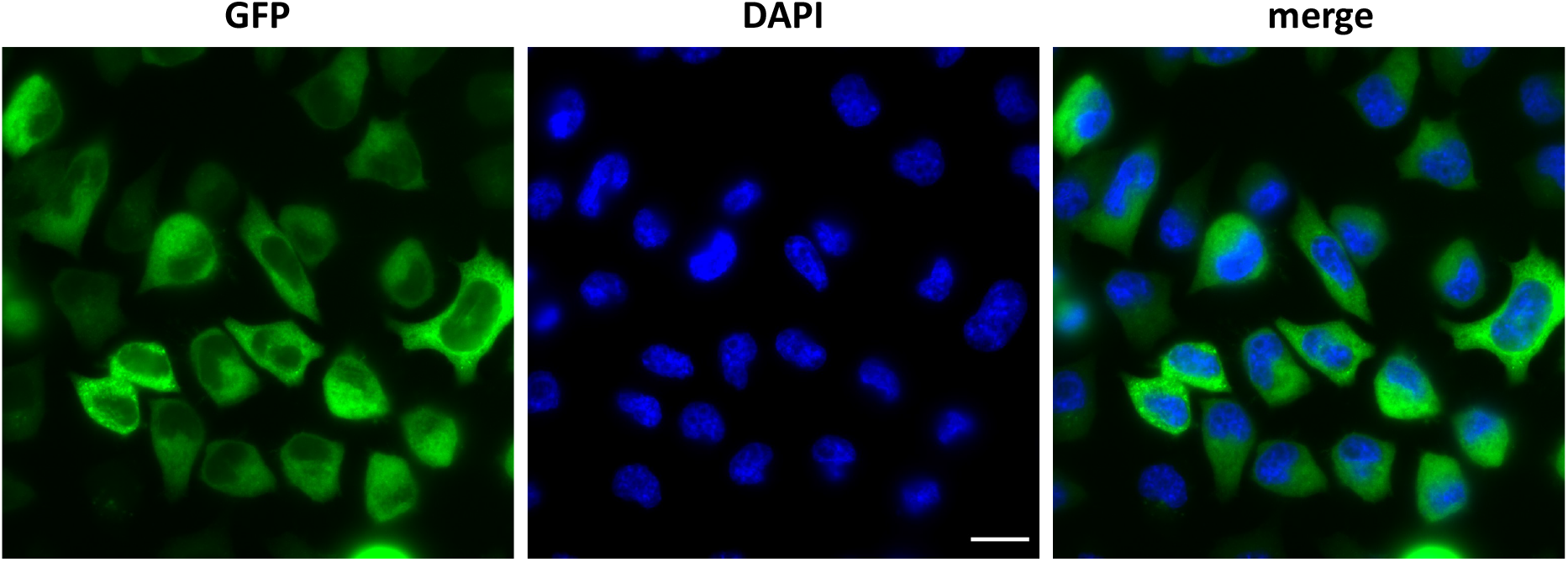
GFP-HELZ2 is mainly localized in the cytoplasm. Representative microscopic image of HeLa cells transfected with a plasmid expressing GFP-HELZ2. Green: GFP signals, blue: DAPI signal, and merged. Scale bar: 15 μm.

## Discussion

HELZ2 appears to be an extremely large and complex protein whose expression is tightly regulated by several mechanisms including induction by interferon and unusual translation initiation. The presence of homologs in many metazoan groups including vertebrates but also invertebrates such as some corals and mollusks suggest that it arose early during the evolution of this group of organisms. It is noteworthy, however, that it is absent from some lineages such as insects, suggesting that it has been lost when organisms encountered conditions in which its function wasn’t advantageous.

The presence of a RNB domain prompted us to analyze the function of the HELZ2 human protein for which biochemical activities were lacking. Careful compilation of available evidence, including transcript mapping data, ribosome profiling results, and evolutionary information suggested that the human HELZ2 protein contained a conserved N-terminal extension not documented in current databases and that its translation initiates at a non-canonical GUG codon. This possibility was confirmed experimentally. In this context, it will be worth reinvestigating whether the previously described HELZ2α and HELZ2β isoforms are expressed and their biological relevance. Translation initiation at GUG codon has been reported previously in human cells for viral mRNAs as well as for specific cellular mRNAs such as DAP5/eIF4G2 (38–40). Indeed, transcriptome-wide ribosome profiling experiments (41) and data mining (42) indicate that GUG is, with CUG, the most common substitute to AUG to initiate translation (reviewed in (29)). In this context, it is noteworthy that expression of HELZ2 (PRIC285) in B cells has been shown to be modulated by a polymorphic locus controlling the level of the RPS26 ribosomal protein (23). Given the role of RPS26 in translation initiation, in particular its interaction with eIF3 and with the mRNA upstream of the start codon (43), it will be of interest to test whether altered RPS26 level modulates HELZ2 translation initiation. Indeed, inefficient initiation at the GUG codon will likely allow recognition of one of the downstream AUG codon, leading to translation of short ORF and probably mRNA degradation by NMD. Taking into account that HELZ2 expression is induced by interferon and the wide impact that this cytokine has on translation (44), future analyses should test whether it modulates the efficiency of translation at non-AUG codon, particularly in the case of HELZ2.

The RNB domain of human HELZ2 contains a substitution of a key aspartate residue in its catalytic center by an asparagine. Yet, our data demonstrate that HELZ2 is an active ribonuclease with characteristics similar to other RNB 3’-5’ exonucleases. The activity of HELZ2 was somewhat unexpected given the conservation of this amino acid in RNB protein. Indeed, this aspartate had been previously proposed to participate to the coordination of catalytic magnesium. However, recent structural analyses reveal that it is involved in interaction with the substrate RNA rather than in the catalytic center (35), a situation that may explain why its substitution with asparagine is compatible with activity. Thus, HELZ2 is the fourth active member of the RNB family in human together with DIS3 and DIS3L1 that are catalytic subunits of the exosome and Dis3L2 a cytoplasmic protein acting on its to degrades RNA substrates harboring a poly-uridylated 3’ end tail (45–47). Like DIS3L2, our data indicate that HELZ2 preferentially degrades substrate ending with U residues. Such substrates are often generated *in vivo* by the action of Terminal Uridyl Transferases (TUTase) to mRNA, small RNA and processing fragments (48). Whether HELZ2 will also target these substrates will have to be determined in the futures. Also, the inefficient degradation of poly(A) tail by HELZ2 may suggest that endogenous mRNA may not be affected the primary target of this factor.

Besides its RNB domain, human HELZ2 contains 6 zinc-fingers of two types that may promote binding to single-stranded and double-stranded RNA. Interestingly, highly related zinc-fingers are present in the ZC3H7A and ZC3H7B/RoXaN factors (Supplementay Fig. 3). Both ZC3H7A and ZC3H7B have been linked to some cancers (49, 50). At the molecular level, ZC3H7B/RoXaN was reported to affect miRNA biogenesis by binding specifically to a sequence located in the apical loop of some pri-miRNA. RoXaN also modulates rotavirus infectivity by interacting with the viral non-structural protein 3 (NSP3) that binds the 3’ end of viral RNA. This binding allows the subsequent recruitment of eIF4G to promote their translation at the detriment of endogenous mRNA (51–53). These observations suggest that likewise, the zinc-fingers of HELZ2 may interact with RNA. HELZ2 also contains 2 helicase domains that are related and belong to the Upf1-like subgroup of the SF1 helicase superfamily. Most members of this subfamily have been shown to interact with RNA and proceed in the 5’ to 3’ direction. Interestingly, the closest relative to the HELZ2 helicase domain 1 and 2 is found in the HELZ protein that has been shown to promote mRNA translation repression and decay (54). Some other members of the Upf1-like subgroup of helicases such as ZNFX1, MOV10 and MOV10L1 interfere with viral infections and/or dispersion of mobile elements (55–57) It is noteworthy that, like HELZ2, ZNFX1 and MOV10 are also interferon-inducible proteins (Fig. 2, (58)). Our biochemical data demonstrate that both helicase domains of HELZ2 are active ATPase stimulated by nucleic acids. Most importantly, they collaborate with the RNB domain, allowing the latter to degrade RNA protected by base-paring with a complementary RNA (Fig. 6), provided that the complementary strand contains a single-stranded 5’-3’ overhang that probably acts as an entry point for one or both helicase domains. Further work will be necessary to understand how the unique combination of domains present in HELZ2 permits the degradation of structured RNA. In this vein, it is of interest to note that mutation in patients suffering from primary biliary cholangitis affect an amino acid located between the RNB domain and the final helicase module. This substitution of a threonine by a methionine may affect the coordination of the different domain and may compromise its activity, contributing to altered immunological reaction responsible for this disease (23, 24).

Based on the study of knock-out mice, HELZ2 inactivation has been described to result in increased leptin receptor mRNA in the liver, leading to a reduced lipogenesis and amplified fatty acid oxidation. Consequently, mutant mice were resistant to high-fat diet-induced obesity, glucose intolerance, and hepatosteatosis (25). The molecular mechanisms by which HELZ2 affect these processes aren’t yet deciphered. In particular, while guanabenz acetate has been reported to bind HELZ2 and limit fatty liver and hyperglycemia associated with obesity, we couldn’t observe an effect of this molecule on the RNase and ATPase activities of HELZ2. The characteristics of HELZ2 including its induction by interferon and nucleic acids, as well as its capacity to degrade structured RNA make it a prime candidate to target viruses and mobile elements that have often compact and structured organization either at the genomic level and/or for their replication intermediates. This is further supported by similarity of regions of HELZ2 with other factors involved in the antiviral response including zinc-finger containing proteins (e.g., RoXaN) and RNA helicases (e.g., MOV10). Consistently, HELZ2 was reported to contribute to cellular defense against Dengue virus (16), HCV (17) and Duck Tembusu virus (21). The cytoplasmic localization of HELZ2 would be consistent with some of these activities. More recently, HELZ2 was also reported to inhibit retrotransposition of human LINE1 (22). In contrast, one report suggest that HELZ2 may have a proviral action for SARS-Cov2 (20). However, this statement should be taken with caution given that the CRISPR strategy used to reach this conclusion may not have taken into account the peculiar structure of the HELZ2 gene with its targeting potentially interfering with non-canonical translation initiation and expression of the extension described above. This issue should be further explored, especially taking into account the reported interaction of HELZ2 with the N protein of SARS-COV2 (59).

Several ribonucleases acting in response to viral infections are already well characterized (for review, 60). For example, the major OAS/RNase L pathway is activated during viral infection by interferons. Once activated, RNase L will degrade exogenous RNA and ribosomal RNA to block virus production (61–63). RNase L will also degrade cellular mRNA involved in mRNA stability and translation, differentiation and interferon-stimulated genes (63). However, several viruses have developed a way to escape degradation by RNase L (64–67). The understanding of the contribution of HELZ2 to the defense against viral infection is then of great interest in the context of the continuous race between viruses and their host.

## Funding

This work was supported by the Ligue Contre le Cancer (Equipe Labellisée 2020) [to B.S.] and Projet IdEx Exploratoire 2021 [to B.S.]. This work of the Interdisciplinary Thematic Institute IMCBio, as part of the ITI 2021-2028 program of the University of Strasbourg, CNRS and Inserm, was supported by IdEx Unistra (ANR-10-IDEX-0002), by SFRI-STRAT’US project (ANR 20-SFRI-0012), and EUR IMCBio (ANR-17-EURE-0023) under the framework of the Plan France 2030.

## Acknowledgments

We acknowledge our team members for support and suggestions. We also thank IGBMC platforms particularly the Imaging and Mediaprep facilities for support.

## Authors contributions

Experimental design, E.H. and B.S.; initial experiments J.S.; experiments realization E.H.; mass spectrometry analyses, B.M.; writing of original draft, E.H. and B.S.; manuscript reviewing, E.H., J.S., B.M., B.S.; funding and project supervision, B.S.

## Declarations of interest

There are no conflicts to declare.

## Notes

### Competing Interest Statement

The authors have declared no competing interest.

