## Supplementary data for "HELZ2: a new, interferon-regulated, human 3’-5’ exoribonuclease of the RNB family is expressed from a non-canonical initiation codon"

Eric Huntzinger[1] (ORCID 0000-0001-6126-4227), Jordan Sinteff[1,2], Bastien Morlet[1] and Bertrand Seraphin[1*] (ORCID 0000-0002-5168-1921)

[2]Present address: 77500 Chelles, France

[*]Corresponding author

**Supplementary Figure 1: Nucleotide sequence of HELZ2 mRNA 5’end.**

exon 1 ┬ exon2

agaaucgaaacugagagcuccugggcaggcucggcagggcaggcagcuccaggagggcuucgaaccguggccaacaguuccag

uggacugcguggacccgugagcucaggagccucagacgccucccuggagagccaagcugguguucgagGUGGCGCCUCCAGGG

UCCACCCUGCUGCCCAACAGCCCCGCGGCCACCAGAGGGCCGUCCCUGGCCCGGCUGUGUGCCCUGGUGGACCUGUGUCUGGG

sORF 1 start — within sORF 1

CUGCUCCCGCUGCACCCAGCGGCUCAAUGAAAGCACCUACGUCCUCCGUAGGGUGGAGCAUGACUGCUCCCGCGAGAUCCUGC

UGGCCCGCUUUAAGCAGGCCACCAAGAGCAAGGUCUGGCGCGUGGUGGGCUGCCGGCCCACCUUCCCAAGGCCCCUGUGCUAC

CAAGUCUGCCACUACUACAGCCCUGGGCUCGGCUGCCGGCGCCACCGAAACCGGUGCACCUUUGCCCGCAGUCGCGAGGAGGC

sORF 2 start

CCUGGUCUGGACCUUCGAGCGUCAGCACAACCUCCAGCGCCUAUGGCUGAAGGCGGAGGUGCAGGGCAGCGGGGCCCAGGGAG

sORF 1 stop

GGGCAGGCCGGGCGGCCGACGCCAUCCUUACGGAGUUUGGCGGCCGCUUCGAGCUGCUUUGCUCCCUCUGCUUCAGGCGCUGU

within sORF 2

CCCCCAUGCAUCUGUCGCGUGGACCCCCAGGGGCAGUGCCCUGAGCACGGAGCCUGCCCCUCCCUCCUGGCCCACGUGAGCGC

sORF 2 stop

CGAGGGCCGCCGCAAGCAACAGUUUGUGGUGGUGAGGCCGCGGCCCCGGGCCGGCCAGCCUCCUGCCUACUGCAGGUUUGUGG

HELZ2β start

GGCGUGGGCAGCCGUGCUGGCGUGGGGAGUCCCGCUGCCAGUUUGCACACAGCGCCGUGGAGAUG

The first 895 nucleotides of HELZ2 mRNA, until the AUG reported to initiate the HELZ2β isoform, are shown. Lowercases are the 5’UTR nucleotides, uppercases are nucleotides of the coding sequence of the N-terminal extension. Dark green boxes highlight proposed non-canonical initiation codons in favourable context. Light green boxes highlight other putative non-canonical initiation codons. Salmon boxes highlight the initiation and stop codons of the 2 sORFs that are also presents and the codon reported to initiate translation of of HELZ2β isoform. Note that both of the two sORFs present in the 5’ UTR contains internal AUG codon that could also promote translation initiation.

**Supplementary Figure 2: Overview of riboseq and ribosome profiling data supporting translation inition at the GUG codon.**

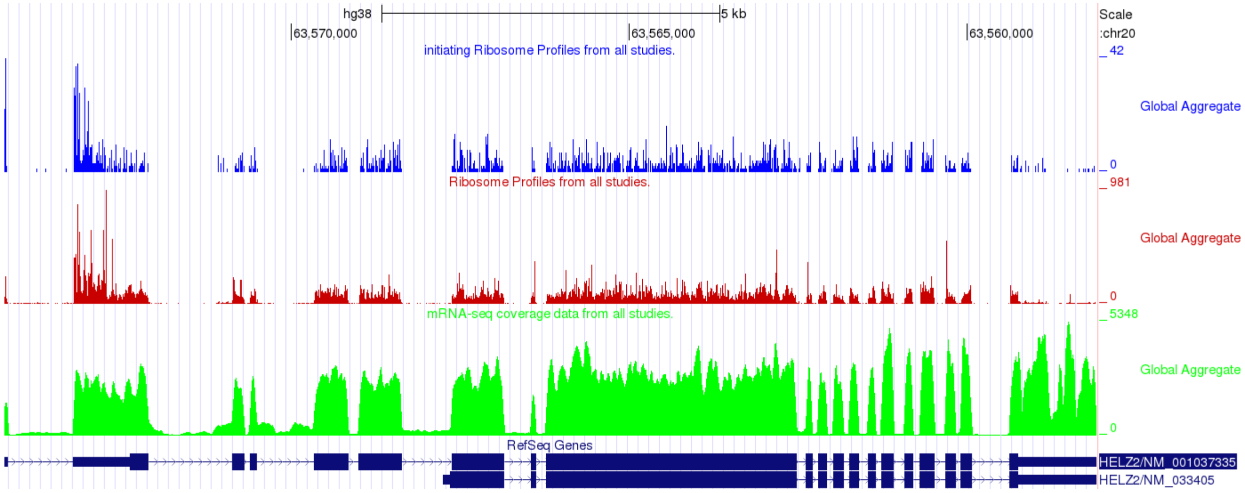


Image obtained from the GWIPS-viz website. Light blue: global aggregate of initiating ribosomes from 8 different studies. Red: global aggregate of ribosomes profiles from 46 different studies. Green: mRNA-seq coverage from 34 different studies. Dark blue: exons-introns organization of annotated mRNAs encoding the HELZ2β and HELZ2α HELZ2 isoforms, respectively. Previously described UTR regions are shown as smaller rectangles.

**Supplementary Figure 3: Alignments of the zing fingers present at the terminus of HELZ2 proteins from various vertebrate species and from the human ZC3H7A and ZC3H7B proteins.**

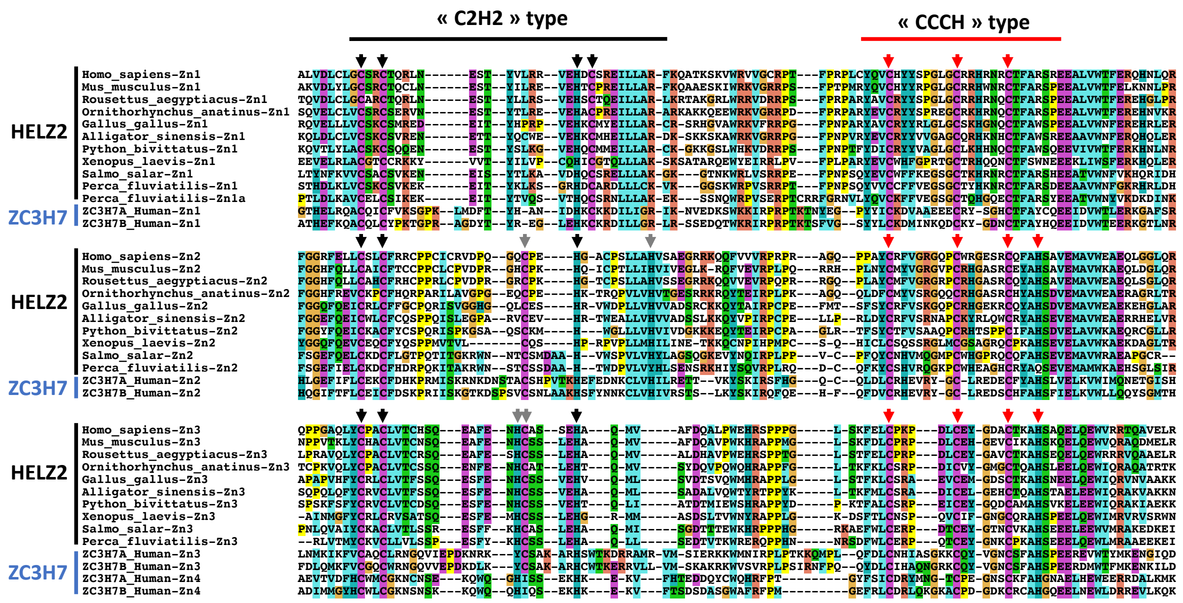

Alignments were produced and coloured for amino acid conservation with Clustal W. Conserved C and H residues likely to be involved in zinc coordination are indicated by black, red and grey arrows. The region corresponding to C2H2 and CCCH zinc fingers and their derivatives are indicated above the sequence. Both for HELZ2 and ZC3H7 proteins, C2H2 and CCCH zinc fingers are alternating. Three tandem copies are usually present in HELZ2 (Zn1-Zn3) and 4 in ZC3H7 proteins but a supplementary copy of 2 zinc fingers in some fish (Zn1a *Perca fluviatilis*) substantiating the importance of this tandem organisation. Note that some zinc fingers are unconventional and possibly non-functional (e.g., the Zn1 CCH finger that lacks a H residue). Some C2H2 fingers also depart from the canonical organization with the presence of potentially alternative coordination residues (grey arrows) in Zn2 and Zn3 fingers. Some of these features being conserved between HELZ2 and ZC3H7 proteins, they may reflect a higher order folding of the region.

**Supplementary Figure 4: Alignments of the full-length sequences of HELZ2 proteins from various vertebrate species.**

### Supplementary Figure 4 (1)

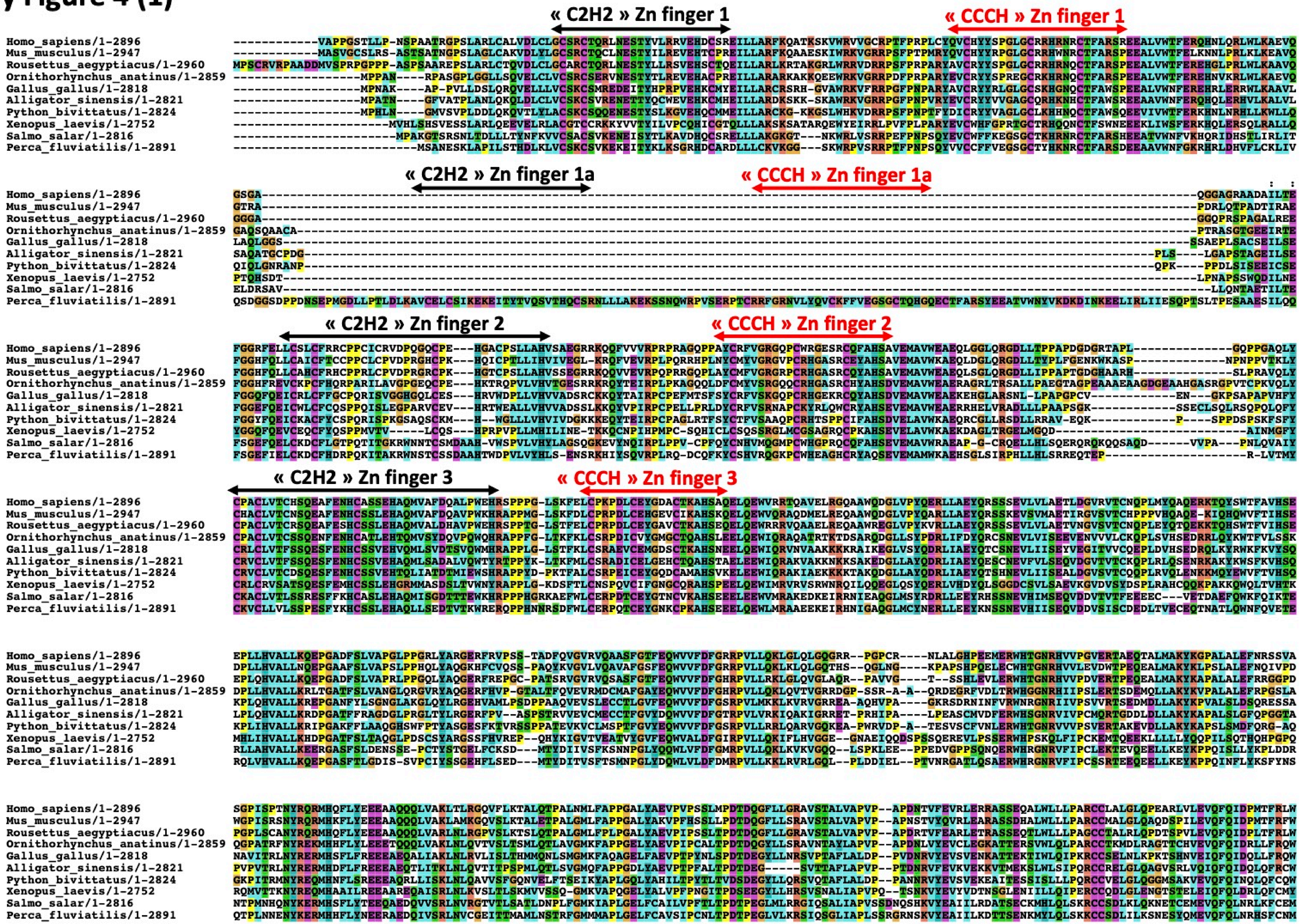

### Supplementary Figure 4 (2)

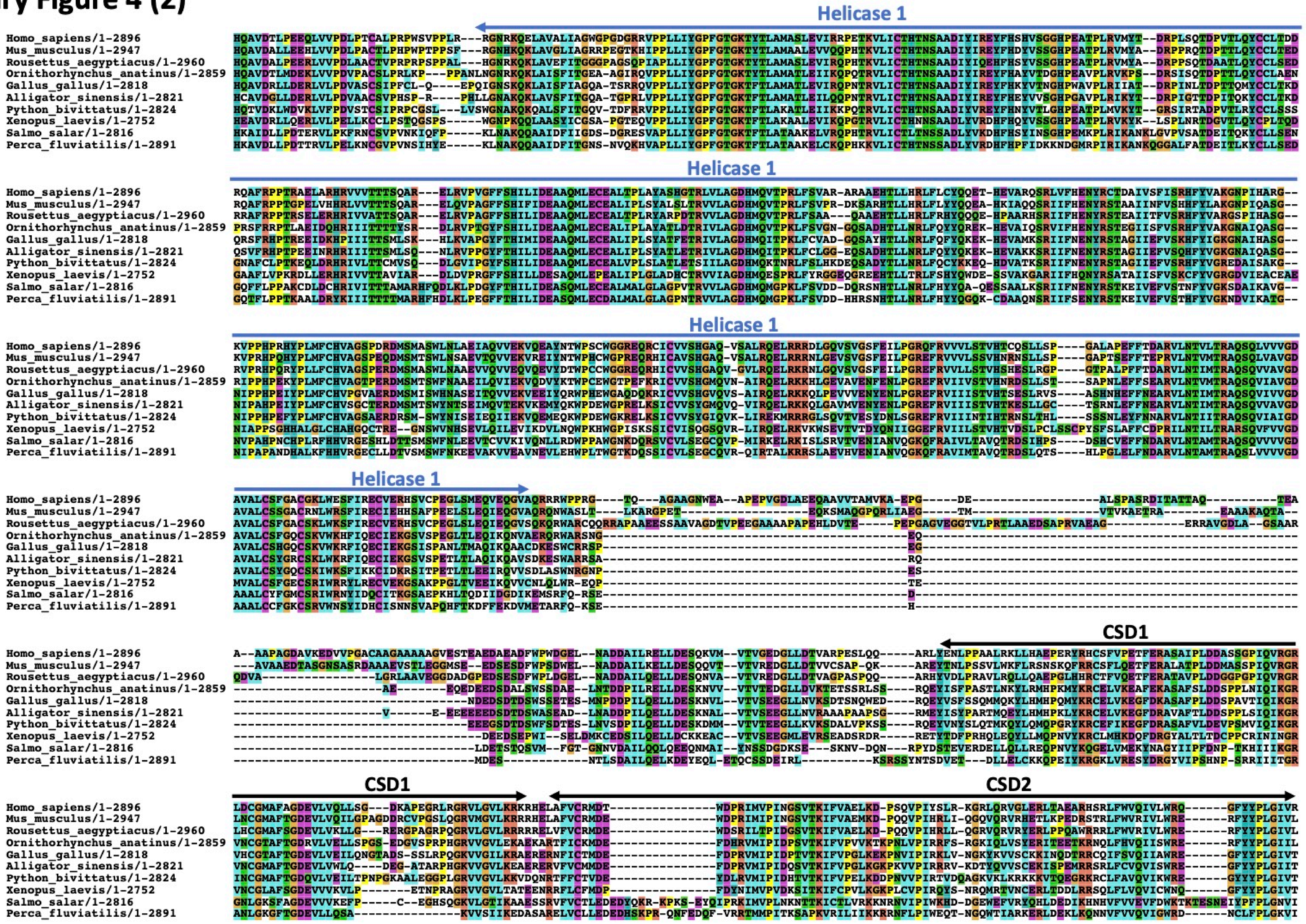

### Supplementary Figure 4 (3)

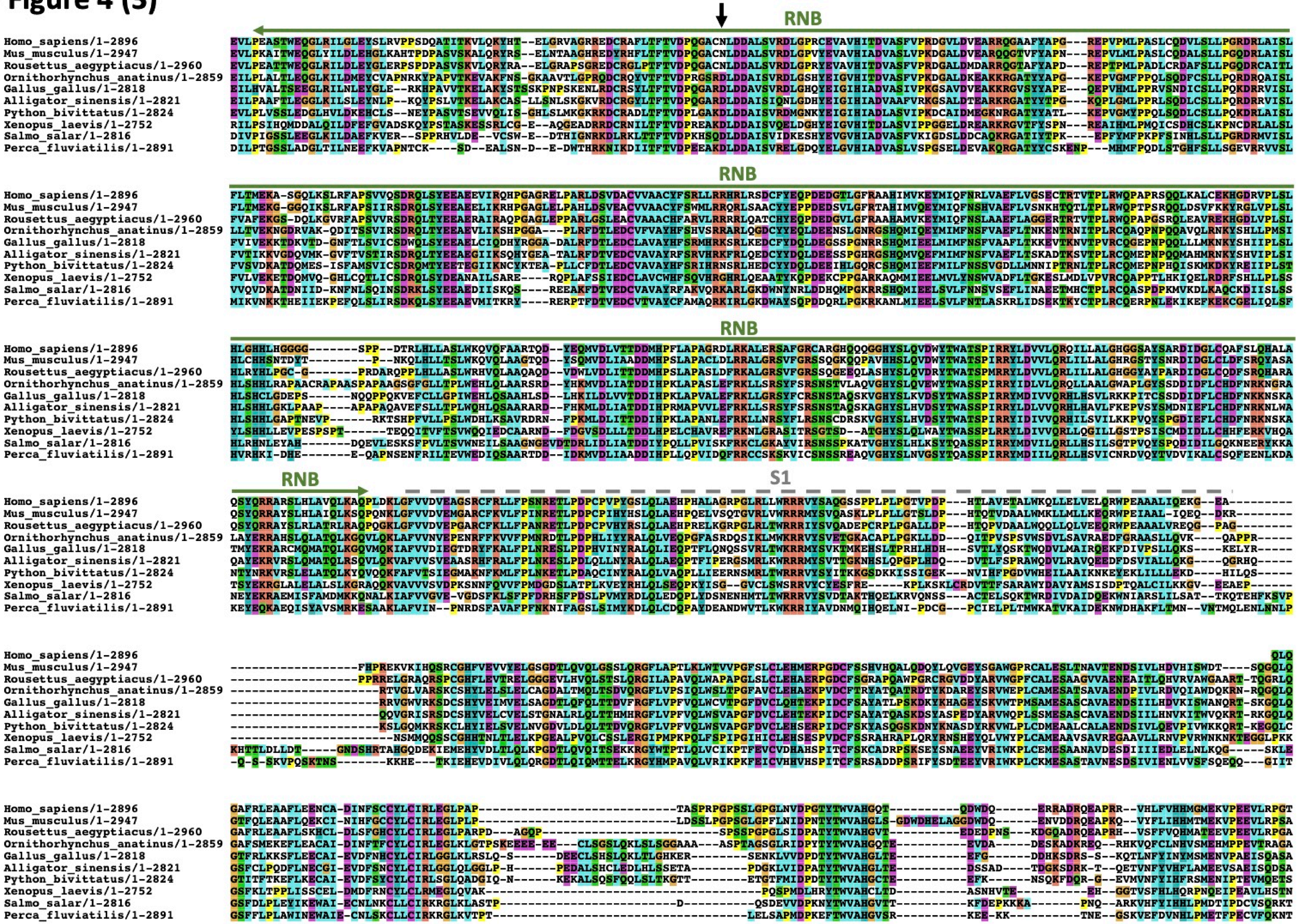

### Supplementary Figure 4 (4)

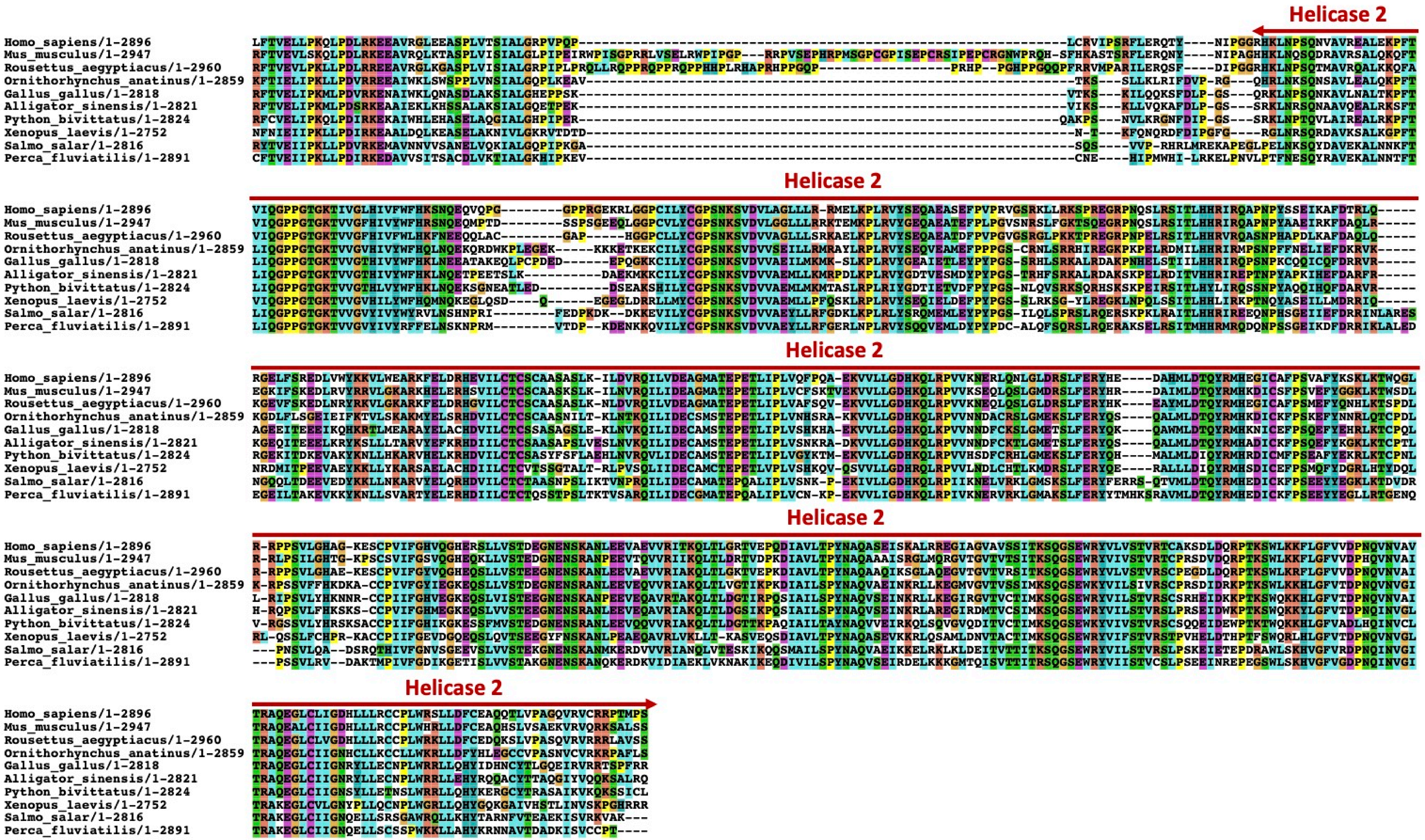

Alignments were produced and coloured for amino acid conservation with Clustal W. The location of various domains with borders are indicated above the sequence. Note that the location of the S1 domain remains to be confirmed. The residue substituted in the catalytic site of HELZ2 from some mammals but conserved as an aspartate in monotremes and other vertebrates in indicated by a black arrow.

**Supplementary Figure 5: HELZ2 and DIS3L1 produce similar final degradation products.**

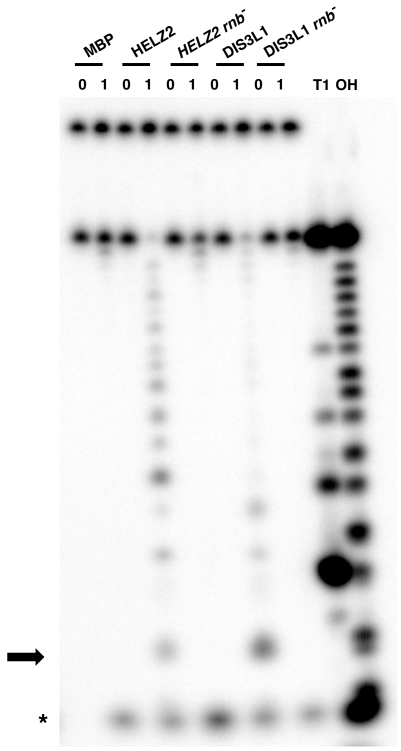


Representative RNA degradation assay of the RNA20 substrate in presence of HELZ2 and DIS3L1 wildtype or mutant thereof carrying catalytic site mutations. Samples from the ribonuclease activity test were fractionated on a 20% acryl-urea 8M gel. Numbers above the lanes represent incubations times in hours. Black arrow: size of the final degradation product. T1: RNase T1 digestion of the substrate. OH: alkaline hydrolysis of the radiolabelled RNA substrate. Asterisk: non-specific product of the reaction.

**Supplementary Figure 6: Guanabenz acetate does not affect ribonuclease or ATPase activities.**

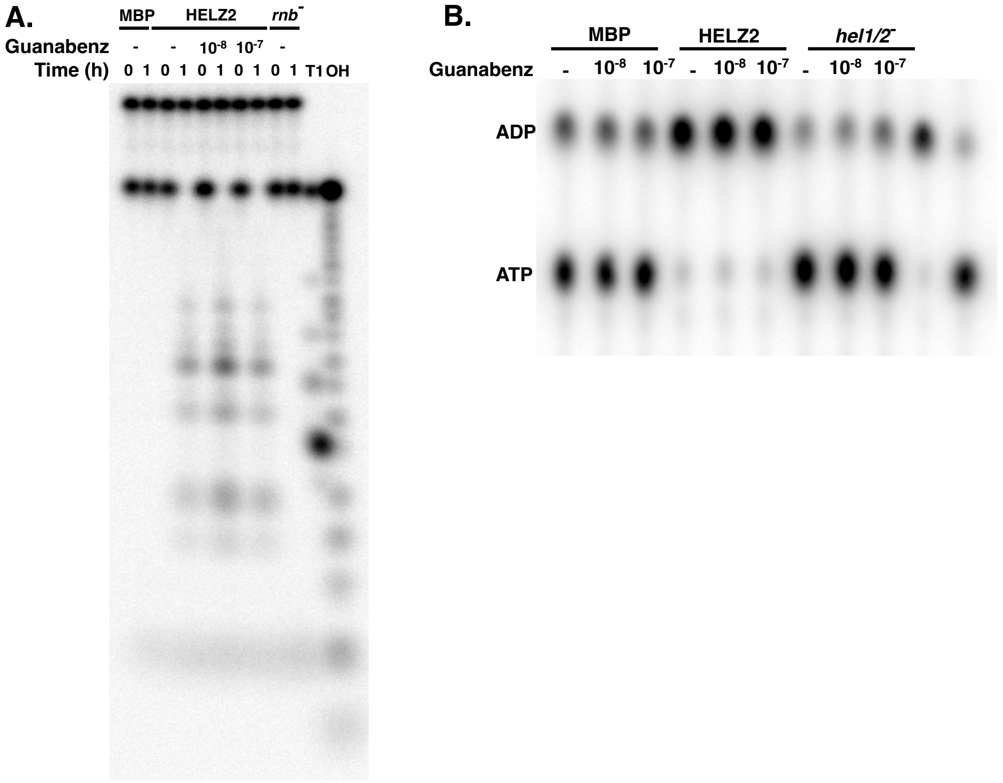


(A) Representative RNA degradation assay of RNA20 substrates in absence or in presence of increasing concentrations of guanabenz acetate. Samples from the ribonuclease activity test were fractionated on a 20% acryl-urea 8M gel. Numbers above the lanes represent incubations times in hours. T1: RNase T1 digestion of the substrate. OH: alkaline hydrolysis of the radiolabelled RNA substrate. Asterisk: non-specific product of the reaction. (B) Representative ATPase assay analysed by thin layer chromatography. Reactions were performed for 2 hours in the presence of ssDNA with the indicated concentrations of guanabenz acetate. The final 2 lanes are markers to identify the distance of migration of ATP and ADP, respectively.
